# A simple mung bean infection model for studying the virulence of *Pseudomonas aeruginosa*

**DOI:** 10.1101/229757

**Authors:** Sneha Garge, Sheyda Azimi, Stephen P. Diggle

**Affiliations:** Department of Microbiology and Biotechnology center, Faculty of Science, The Maharaja Sayajirao University of Baroda, Vadodara, India; School of Biological Sciences, Georgia Institute of Technology, Atlanta, GA 30032, USA

**Keywords:** Mung bean, *Pseudomonas aeruginosa*, virulence model, quorum sensing

## Abstract

Here we highlight the development of a simple and high throughput mung bean model to study virulence in the opportunistic pathogen *Pseudomonas aeruginosa*. The model is easy to setup and infection and virulence can be monitored for up to 10 days. In a first test of the model, we found that mung bean seedlings infected with PAO1 showed poor development of roots and high mortality rates compared to un-infected controls. We also found that a quorum sensing (QS) mutant was significantly less virulent when compared with the PAO1 wild type. Our work introduces a new tool for studying virulence in *P. aeruginosa*, that will allow for high throughput virulence studies of mutants, and for testing the *in vivo* efficacy of new therapies at a time when new antimicrobial drugs are desperately needed.

## Main text

*Pseudomonas aeruginosa* is an opportunistic pathogen that can be isolated from diverse habitats including water, soil, animals and plants (1-3). In humans, it is a leading cause of infection, morbidity and mortality in cystic fibrosis (CF) lungs, and is a problematic pathogen in burn wounds, chronic diabetic wounds and immunocompromised individuals (1, 3, 4). Because of the problems that *P. aeruginosa* causes in human hosts, researchers have developed a number of different *in vivo* infection models for assessing *P. aeruginosa* virulence, pathogenesis and disease (5-7). These include animal models such as waxmoths (8) (9), fruit flies (10), nematodes, mice (11-13) (14-16), pigs (17) and *ex vivo* pig lungs (18, 19). Because *P. aeruginosa* uses a number of the same virulence factors to infect and cause disease in both animals and plants, plant infection models such as the *Arabidopsis* model have also been widely used (20, 21). A number of factors are important to consider when designing and using a virulence model. These include cost effectiveness, ethical considerations, ease of growth in a laboratory, measures of virulence, and the ability to handle large sample sizes. In this study we describe the development of a mung bean infection model (22), and describe how this easy to establish model to be used for *P. aeruginosa* to assess virulence.

We purchased mung bean seeds from general supermarket stores. To avoid any fungal or other bacterial contamination, we first surface sterilized the seeds by soaking in 70% (v/v) ethanol in a sterile flask for 30 seconds and then we soaked them in 0.01% (w/v) Mercury Chloride (HgCl2) for 30 seconds three times. We then washed the seeds with sterile water. We transferred surface sterilized seeds onto soft agar plates (0.8% w/v agar), and added 2 ml of sterile water to the surface of the agar (Fig. 1, a). We then incubated the plates for 24 hours at 37°C under humidified conditions to allow for the germination of seeds. We used 8 germinated seeds per group for the infection studies. As a first test of our model, we wanted to determine whether quorum sensing (QS) is important for virulence in mung beans, because QS has previously been shown to be an important regulator of virulence in a number of different hosts including mice, *Arabidopsis*, lettuce, nematodes, and insects (23, 24) (11, 25). We used a PAO1 *lasI* ^*-*^ *rhlI*^*-*^ double mutant (PAO-JG1) (26) to determine the role of QS on pathogenesis, colonisation and disease development in mung beans.

**Fig. 1.**
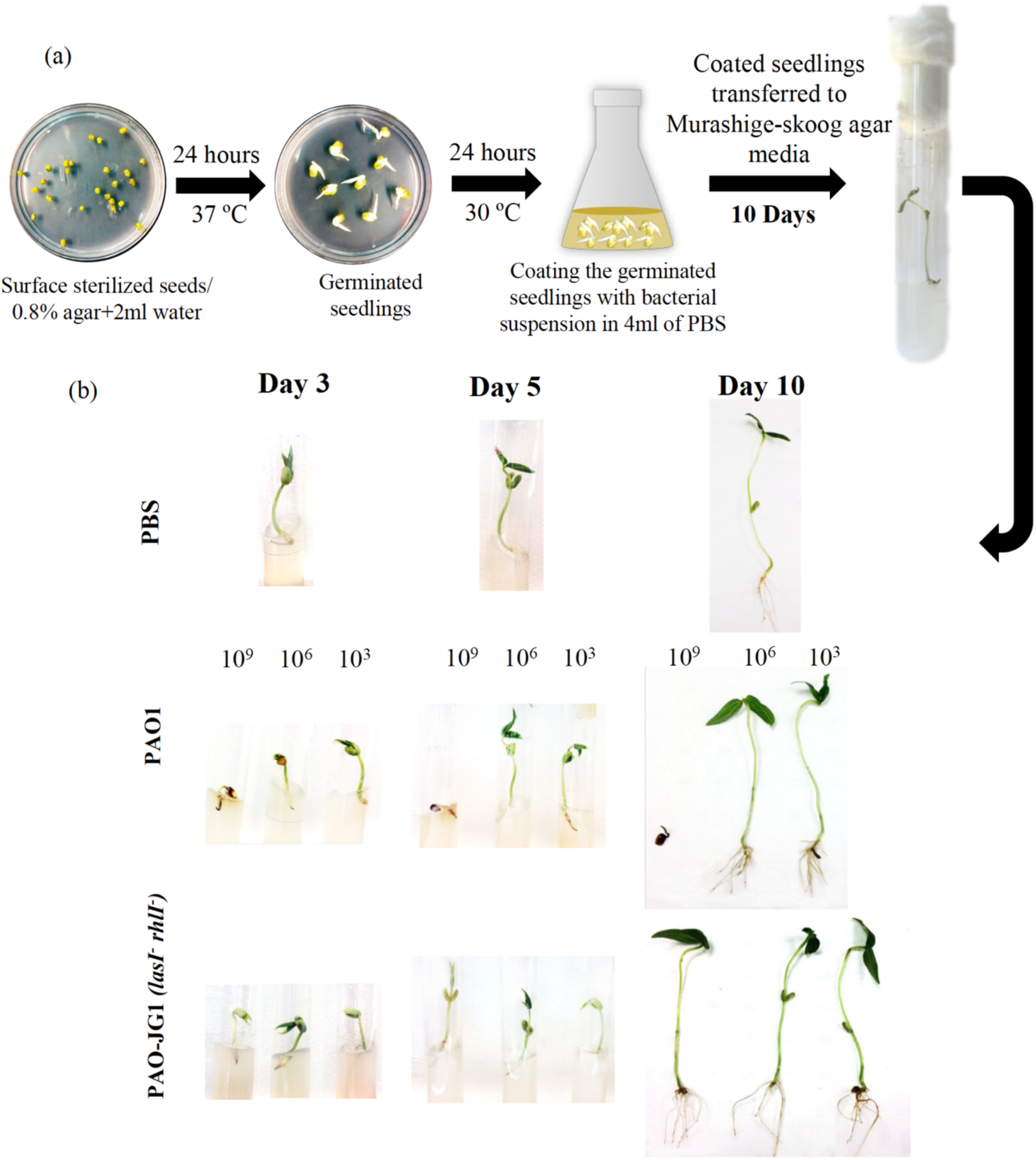
Mung bean infection method and comparison of growth defect in infected plants. (a) The procedure to infect mung bean seeds with *P. aeruginosa*. (b) the comparison of PAO1 and PAO-JG1 treated mung bean plants with different initial bacterial densities after 3, 5 and 10 days post treatment (images are representative of 3 independent trials).

To infect the seeds, we inoculated the PAO1 wildtype strain and PAO-JG1 into 100 ml Lysogeny Broth (LB) from overnight start-up cultures, and incubated at 30°C/200 rpm to reach a cell density of ∼10^9^, 10^6^ and 10^3^ Colony Forming Units /ml (CFU/ml). We centrifuged stationary phase cultures at 7000 rpm for 10 min and we re-suspended pellets into 4 ml of sterile Phosphate Buffered Saline (PBS) (pH 7). We added 8 geminated mung bean seeds to the bacterial suspension and then incubated at 30°C for 24 h under static conditions to achieve a good colonization of bacterial cells on the germinated seeds. We incubated equal numbers of seeds in 4 ml of PBS as uninfected controls. After the incubation, we took 3 seeds from each treatment group, and re-suspended them in 1 ml of sterile PBS and vortexed for 30 s. We used this suspension to perform viable counts (CFU/ml), to determine the number of adhered bacteria that coated each seed. In our first trial runs, we used all three initial CFU/ml of bacterial cultures, however we saw no significant differences in growth rate of plants infected with PAO1 wildtype less than ∼10^9^ CFU/ml (Fig. 1, b).The cell counts recorded are shown in Table 1.

**Table 1.**
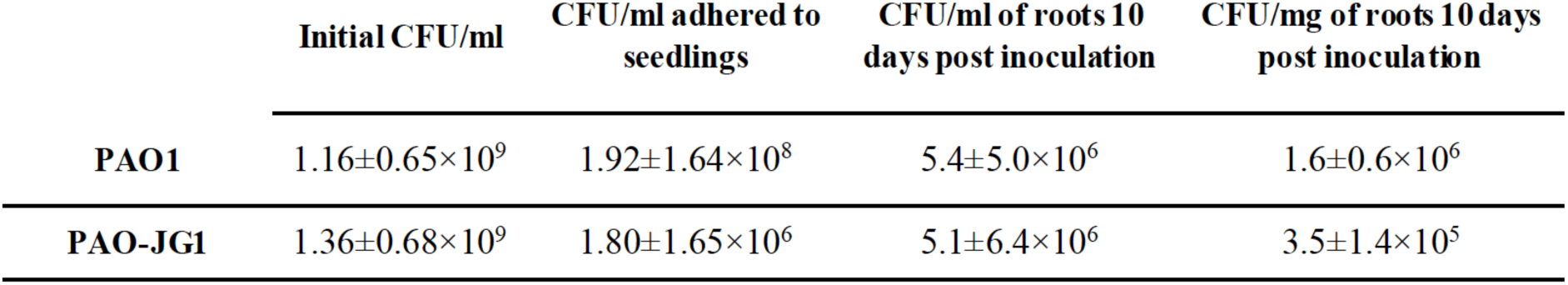
Viable counts of colonized bacteria in seedlings and roots by PAO1 or PAO-JG1. Values are represented as mean± SD for six independent trials with five technical replicates.

|  | Initial CFU/ml | CFU/ml adhered to seedlings | CFU/ml of roots 10 days post inoculation | CFU/mg of roots 10 days post inoculation |
| --- | --- | --- | --- | --- |
| **PAO1** | $1.16 \pm 0.65 \times 10^9$ | $1.92 \pm 1.64 \times 10^8$ | $5.4 \pm 5.0 \times 10^6$ | $1.6 \pm 0.6 \times 10^6$ |
| **PAO-JG1** | $1.36 \pm 0.68 \times 10^9$ | $1.80 \pm 1.65 \times 10^6$ | $5.1 \pm 6.4 \times 10^6$ | $3.5 \pm 1.4 \times 10^5$ |

For plant growth, we prepared Murashige-skoog agar media [Murashige-skoog 4.4 g/L, (Sigma aldrich) and 0.8% w/v agar (Oxoid)]. We poured 30 ml aliquots of media into large glass tubes (30×200 mm, Duran groups) and sterilized by autoclaving for 20 min. We aseptically transferred both infected and PBS control mung bean seedlings (5 seeds per group) into the tubes. We allowed the seedlings to grow in a natural daylight cycle for 10 days in gnotobiotic conditions. To determine viable bacterial counts (CFU/mg of root weight) of both the strains colonizing the roots of the mung bean seeds during the incubation of 10 days, we cut the root of each plant with sterile scissors, and re-suspended these in 1 ml of sterile PBS and vortexed for 30 s. We used this suspension to perform a viable count of bacteria to determine the number of bacteria colonizing the root of each plant. We normalized the CFUs on each root by the respective root weight (mg) of each plant root. We show that both PAO1 and PAO-JG1 strains were viable on the roots throughout the infection period (Table 1). We determined the health of the plants by measuring parameters such as mortality rate of seedlings, shoot length, root length and the number of root branches due to the infection. We defined mortality rate as the percentage of seedlings that died from the total number of seedlings treated with bacterial culture suspension in each trial. We found that the mortality rate of the seedlings treated with PAO1 was higher compared to PAO-JG1 treated seedlings (Table 2). We observed all the seedlings infected with various CFU/ml of PAO-JG1 developed into healthy plants (Fig. 2, a and Fig.1 b). We found that seedlings which survived infection with approximately 10^9^ CFU/ml of PAO1, demonstrated severely attenuated growth (Fig. 2, a and Fig. 1, b). These plants displayed disease symptoms such as water-soaked lesions on their roots and blackening of the root and shoot.

**Fig. 2.**
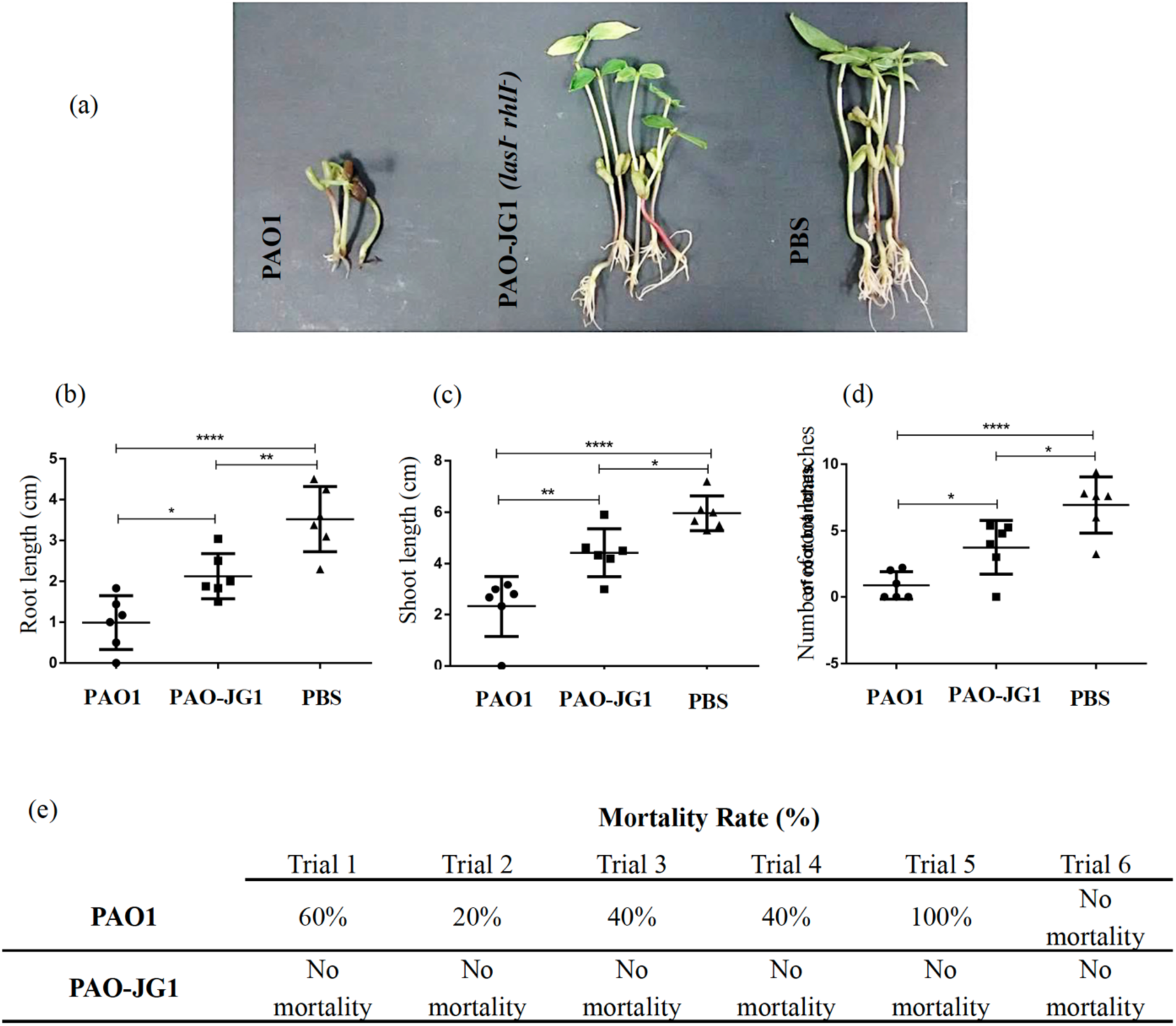
Mung beans infected with QS mutant strain PAO-JG1 grow significantly better that seed infected with PAO1. (a) The representative of PAO1 and PAO-JG1 treated plants (b) root length; (c) shoot length and (d) number of root branches. Values represent the mean of six trials. Bars indicate standard deviation of the mean. Each trial has five replicates. Statistical analysis was done using one-way ANOVA Bonferroni’s multiple comparison test (* = *p<*0.05, ** = *p*< 0.01 and *** = *p*< 0.0001). (e) Mortality rates in mung bean treated seedlings by PAO1 and PAO-JG1. There were five culture suspension treated seedlings in each trial.

Our infection model examined three parameters to determine and score plant health: root length, shoot length and the number of root branches. In seedlings that survived PAO1 treatment, we observed a significant decrease in the root and shoot length measures when compared to seedlings treated with PAO-JG1 and healthy PBS control plants (Fig. 2, b, c and d). We found that the root development of PAO1 treated plants was also impaired as the number of root branches were reduced when compared to PAO-JG1 treated plants and healthy PBS control plants (Fig. 2, d).

In 6 independent trials, each with 5 technical replicates, we found that PAO1 either caused the death of the seedlings or the seedlings survived with impaired growth. In contrast, we found that PAO-JG1 did not cause mortality of seedlings in any of the trials and furthermore, the seedlings developed into significantly healthy plants after 10 days (Fig. 1 and Table 2). These data demonstrate previous observations that the *las* and *rhl* QS systems regulate the virulence of *P. aeruginosa* in plant tissues (27-29). Although we found that QS is important for virulence and disease in mung beans, infections were not completely prevented, suggesting the role of other factors responsible for pathogenesis in mung bean plants which are QS-independent.

Our data demonstrates that we have developed a simple plant model to study mechanisms of pathogenesis and virulence during *P. aeruginosa* infection. Our model has advantages compared to the popular *Arabidopsis* model, which takes 4-6 weeks to grow the plants, and then 5 days post inoculation for observing disease symptoms (20, 27). The mung bean model takes a maximum of 12 days, which includes 1 day to obtain germinated mung bean seedlings, 1 day for infecting the germinated seeds of mung bean with bacterial cells and 7-10 days to grow the plants and allow for the development of disease symptoms. The ease of the methods involved in this model also offers a number of conveniences. It is easy and cheap to acquire mung bean seeds, germinate them and maintain the plants in most lab facilities and without any ethical considerations. Coating germinated seedlings with culture suspension to achieve infection is a simple microbiological technique, and coating allows researchers to evaluate infection in terms of colonization and mortality of seedlings. This can be used as an important parameter for qualitative studies of the colonization of a host, virulence and phenotypic traits involved in the pathogenesis of *P. aeruginosa*. You can also score the pathogenesis of *P. aeruginosa* in plants by measuring parameters such as root length, shoot length and the number of root branches. These are useful parameters where levels of infection by different mutants or strain can reveal important information about virulence factors and virulence in general.

Large sample sizes is not a limitation for the model, so it can be used as a high-throughput virulence screen. Due to the clear effects that we found of QS on virulence; screening libraries of chemical compounds such as *N*-acyl homoserine lactone (AHL) mimics, analogues, inhibitors, antagonists, or quorum quenching bacteria for controlling the infection, could be easily tested. Functional studies of the genes and regulatory proteins which are involved in phenotypes such as motility, adhesion, colonisation, pathogen survival, and their impact on virulence can be conducted using the model. Finally, if seedlings are coated with sub-lethal doses of *P. aeruginosa*, which do not cause the death of plants, this could allow for sociomicrobiology studies relating to social behaviours, to determine how spatial and temporal interactions within and between bacterial communities progress in a natural environment. In summary, we have developed a cheap, reproducible and simple to cultivate plant host model which can be used to study *P. aeruginosa* virulence, and it also may also have relevance for other bacterial species.

## Acknowledgments

S.G. was sponsored by an EMBO Short Term Fellowship (ASTF number: 27-2015). This work was supported by a Human Frontier Science Program Young Investigators grant to S.P.D. (RGY0081/2012). We thank Freya Harrison for helpful comments on the work.

We have no conflict of interest to declare.

